# Chemical mate choice copying in *Drosophila melanogaster*

**DOI:** 10.1101/2024.06.23.600307

**Authors:** Bloo Mitchell, Alexandria Abbott, Ashanti Brown, Lacy Skinner, Elmira Umarova, Philip Kohlmeier

## Abstract

Mate choice is a critical decision especially for females that requires time and energy to assess potential partners’ genetic quality. Consequently, in many species, females have evolved the ability to utilize social information by copying the mate choices of others, usually based on visual cues. However, many species, especially invertebrates, primarily rely on chemical not visual cues. Using chemical rather than visual cues provides several advantages such as not requiring active observation of copulations. Despite of that, empirical evidence for the existence of chemical mate choice copying is scarce. Using *Drosophila melanogaster*, we provide the first demonstration of chemical mate choice copying. Females exposed to a recently mated select the same male genotype as the teacher female mated with at a higher frequency than expected by chance. Chemical mate choice copying requires sensing both male and female cues, which might indicate that other females have chosen that male genotype. Our work suggests that females, in the presence of mated females, increase choosiness at the virgin stage, elevating sexual selection on male traits. This study provides novel evidence that exploiting social information is more prevalent in flies than previously assumed.

## Introduction

Selecting a mating partner is a critical decision with significant fitness consequences, particularly for females, as reproduction-associated costs are typically higher in females than in males (Bateman 1948; Trivers 1972; Milinski and Bakker 1992). As a result, females have developed numerous mechanisms to evaluate the genetic quality of potential mates to ensure the acquisition of high-quality sperm. Male traits assessed by females, for instance, include coloration (Hill 2002), courtship song (Champagnon and Cueva del Castillo 2008), courtship intensity (Lin et al. 2016), and pheromonal profiles (Billeter and Levine 2013).

Collecting private information about the fitness of each male can be time and energy-intensive, especially if group size is large and males are abundant (Witte 2007). As an alternative strategy, females across species evolved the ability to exploit publicly available social information by monitoring, memorizing, and integrating mate choice decisions of other females. Such mate choice copying behaviors, whereby a female is more likely to select a male after observing this or a similar male being selected by another female, are widespread and have been documented across taxa including in birds (Drullion and Dubois 2008), fish (Dugatkin and Godin 1993), mammals (Kavaliers et al. 2017), and invertebrates (Mery et al. 2009). Most of the studies that reported the existence of mate choice copying relied on experimental setups that tested for mate copying via visual cues, even if the used model organism primarily communicates via other sensory modalities, such as chemical cues (Scauzillo and Ferkin 2019). This is surprising as olfactory or gustatory cues can convey individual information about the male, such as age (Scauzillo and Ferkin 2019), dominance status (Scauzillo and Ferkin 2019), or parasite load (Claudio-Piedras et al. 2021) that can be relevant for female mate choice. Moreover, females that include chemical cues in their mate copying decisions do not have to be present or actively observe copulations of other females as residuals of the males can still be present on the female after mating. Unlike chemical mate choice copying (CMCC), visual mate choice copying might also be restricted to species that mate during the day due to their dependency on light conditions. Despite the obvious advantage of CMCC, to our knowledge, empirical evidence for the existence of CMCC is lacking.

We tested whether the fruit fly *Drosophila melanogaster* uses CMCC. Fruit flies are social and form mixed sex groups in which multiple males are available as potential mating partners (Billeter et al. 2024). Female flies assess a large variety of phenotypic traits of males during mate choice decisions. These traits include body size (Markow 1988), age (Lin et al. 2016; Zhang et al. 2021), pheromonal profiles (Billeter and Levine 2013), courtship (Villella and Hall 2008), nutritional status (Fricke et al. 2008), and symmetry (Møller and Pomiankowski 1993), among others which might be costly to collect individually. *D. melanogaster* is one of the very few invertebrate species in which mate choice copying has been reported. Female flies that observe another female mating with a poor-condition male are more likely to mate with a poor-condition male themselves compared to naïve females that did not observe any matings before (Mery et al. 2009). This effect was also observable when two similar males dusted with differently colored fluorescent powder were used (Mery et al. 2009; Dagaeff et al. 2016; Danchin et al. 2018; Nöbel et al. 2022*a*), demonstrating that observer females can monitor and use neutral publicly available information and not differences in male condition or attractiveness for mate copying. Females copy mate choices even if the selected male has visible genetic defects, suggesting that social information can overwrite privately collected information (Nöbel et al., 2018). However, *D. melanogaster*, like most other insects, primarily communicates via chemical cues (Wyatt 2003), and Cuticular Hydrocarbon (CHC) profiles are important modulators of attractiveness and mate choice (Billeter and Levine 2013; Kohlmeier et al. 2021). CHC profiles display stable quantitative differences between different *D. melanogaster* strains which are genetically encoded (Jallon 1984; Sureau and Ferveur 1999; Grillet et al. 2006; Billeter et al. 2009). Several male pheromones involved in female mate choice have been identified and include *cis*-Vaccenyl Acetate, Palmitoleic acid, and 7-Tricosene as well as its oxidation product heptanal (Grillet et al. 2006; Kurtovic et al. 2007; Billeter et al. 2009; Kohlmeier et al. 2021; Verschut et al. 2023; Doubovetzky et al. 2024). Some of these pheromones are transferred from the male to the female during copulation and hence, can potentially be sensed by other females in the context of CMCC (Mane et al. 1983; Zawistowski and Richmond 1986; Everaerts et al. 2010; Laturney and Billeter 2016). Despite this, it remains unknown whether *D. melanogaster* or any other invertebrate species uses CMCC.

We used an experimental paradigm in which student females were first exposed to recently mated females and then tested whether or not they had selected the same male genotype as the teacher. Many studies on female mate choice selected male genotypes that do not co-occur in the same environment and potentially overpronounce variation in CHC profiles (Kohlmeier et al. 2021; Doubovetzky et al. 2024) which questions to what extent the studied behaviors are ecologically relevant. We circumvented this issue by using two isofemale lines generated from flies collected in the same environment and, thus, represent naturally occurring variation in male traits.

## Material and Methods

*D. melanogaster* flies were collected using banana traps in a residential area in Memphis, TN, USA, in July 2023. Two isofemale lines, named *Memphis-4* (*M4*) and *Memphis-7* (*M7*), were generated by transferring two females individually into standard vials (25x 95mm) filled with 10ml fly food following the recipe described in (Kohlmeier et al. 2021) at +25°C and 12:12 Light:Dark photoperiod. Lines were kept in the lab for more than 20 generations before the onset of the experiments.

To test whether *D. melanogaster* females utilize chemical cues in mate choice copying, males from the *M4* and *M7* isofemale lines and virgin females of the laboratory strain *Oregon-R* (*OR*) were aged three to seven days in groups of 15. In contrast to previous studies, we have adopted the terms “student” and “teacher” rather than “observer” and “demonstrator,” as we consider these terms to be more appropriate for the sensory modality examined. Student females were collected without CO_2_ anesthesia to prevent potential effects on memory formation (Muria et al. 2021). These student females underwent one of four treatments (Figure 1):

**Figure 1:**
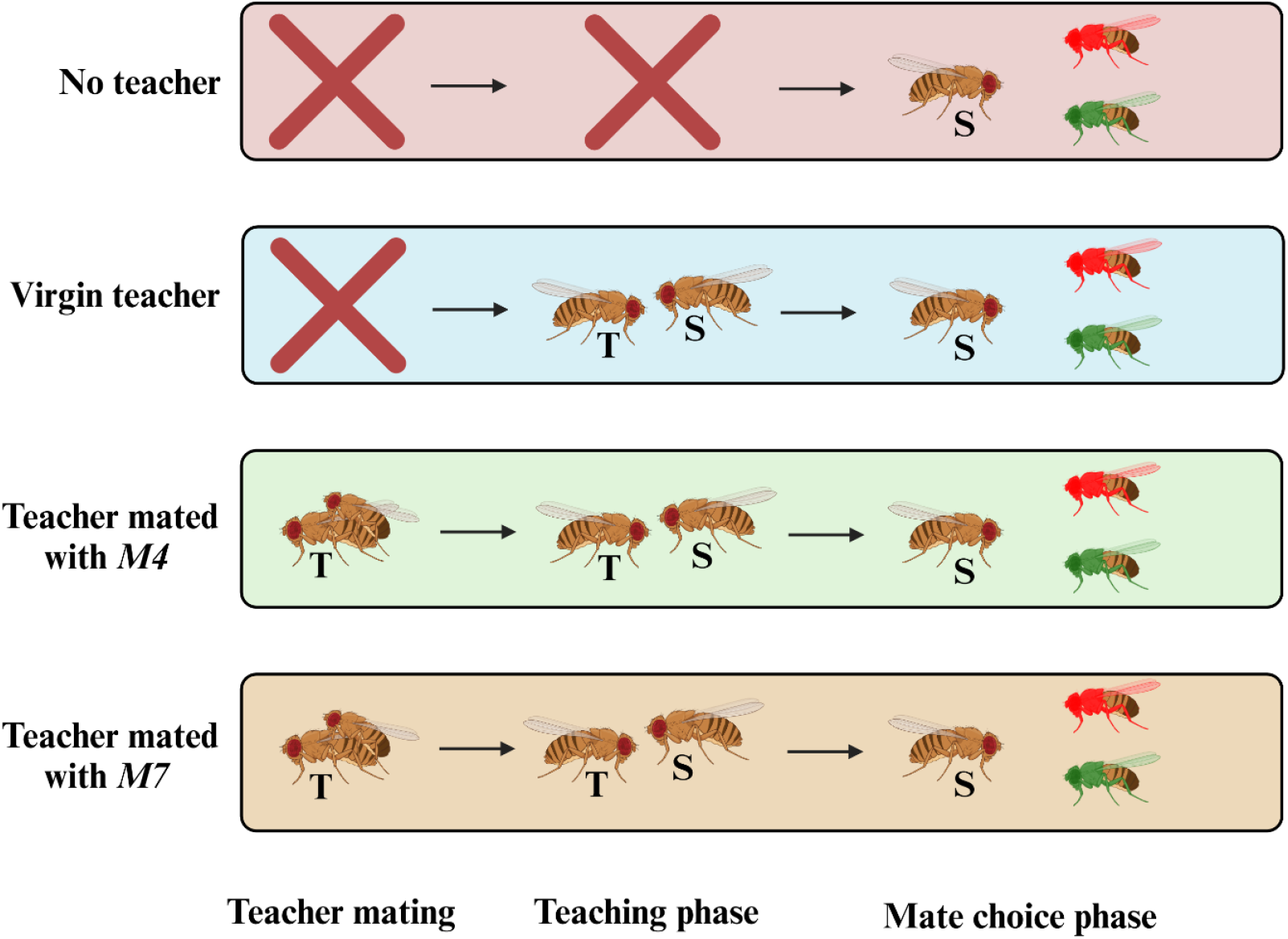
Experimental setup of the first experiment. Teacher females were mated with either *M4* or *M7* males. After mating ended, teacher females were transferred to student females for a 45min teaching phase. Student females were then offered one colored *M4* and one colored *M7* male for mate choice. In the “Virgin teacher” treatment, virgin teachers that had no prior contact with *M4* or *M7* males. In the “No teacher” treatment, mate choice of student females was tested without any teaching phase. T: Teacher. S: Student. Figure created using BioRender.

### No Teacher Treatment

Each female was individually placed into a Greiner Bio-One arena (35mm diameter) containing fly food and one male from each of the *M4* and *M7* lines. To distinguish between the males, they were dusted with pink or green fluorescent powder (Slice of Moon™) at least 30 minutes prior to the experiment following established protocols (Verschut et al. 2022). Females can distinguish even nuanced chemical profile differences between males treated with different dusts (Kohlmeier et al., 2021). Colors for the males were assigned randomly in each trial. Mate choice was recorded, and females who did not mate within 60 minutes were excluded.

### Virgin Teacher Treatment

Two *OR* virgin females collected without CO_2_ were kept together in an arena for 45 minutes. Afterward, one female was randomly selected and transferred to a new arena with one colored *M4* and one colored *M7* male to assess mate choice.

### M4-Mated Teacher and M7-Mated Teacher Treatments

Each *OR* virgin teacher was paired with two males of either the *M4* or *M7* line. At this stage, all males were unmarked by fluorescent powder. After copulation ended, the teacher female was paired in a new arena with a single *OR* virgin student female for a 45-minute “teaching” phase. Afterwards, student females were identified by the absence of a fluorescent mating plug (Lung and Wolfner 2001) using a 395nm Ultraviolet Flashlight (Morplot) and were then transferred to a fresh arena containing one randomly colored *M4* and one randomly colored *M7* male to record mate choice.

We observed that students prefer the male the teacher was mated with. This observation could result from mate choice copying or a “habituation-based preference”, where exposure to a male’s odor – via pheromonal residues left on the teacher after mating – increases the female’s preference for that male. To differentiate between these possibilities, student females were exposed to CHC extracts from *M4*- and *M7*-mated teachers and *M4* and *M7* males, respectively. To prepare these extracts, 15 *OR* virgin females and 15 *M4* or *M7* males were placed together in a vial for 60 minutes to mate. After mating, the flies were immobilized on ice, and mated females were identified by the presence of a mating plug. Three *M4-* or *M7-*mated females or three virgin *M4* or *M7* males that had never encountered a female were then transferred to a 2ml Silanized glass vial (Thermo Scientific). CHCs were extracted by adding 150μl of hexane and vortexing the vial for 3 minutes (Verschut et al. 2022). The extracts were then transferred onto a Whatman filter paper disc (5mm diameter), and the hexane was allowed to evaporate for 30 minutes. The prepared discs were placed in a mating arena, and a student female was added for a 45-minute exposure period. After that, mate choice between *M4* and *M7* males was assessed as previously described.

One prediction derived from the hypothesis that females utilize CMCC to identify attractive males is that when exposed to two teachers, one mated with *M4* and one with *M7*, students should afterward mate randomly as they would have developed a preference for both male genotypes. To test this, *OR* virgin teachers were paired with two males from either the *M4* or *M7* line (also unmarked during this phase). After mating, one *M4*-mated and one *M7*-mated teacher were introduced simultaneously to the student for the 45-minute teaching phase. Following this, the student was placed in a fresh arena containing one randomly colored *M4* and one randomly colored *M7* male to record mate choice.

In the analysis, mate choice outcomes were organized into two columns: one for *M4* males and one for *M7* males, where ‘1’ indicates that the respective male was selected for mating whereas ‘0’ indicates that this male was not selected. Both columns were combined using the *cbind*() function in R to create a matrix of responses. One Generalized Linear Model (GLM) with a binomial distribution was used for each experiment. Each GLM included the outcomes of the mate choice tests as a response variable and treatment (*e*.*g*., mated female extract vs. male extract) as an explanatory factor. In addition, one binomial test per treatment has been used to test whether the preference for *M4* males differs from 0.5, *i*.*e*., random choice.

## Results

The proportion of students selecting *M4* males was influenced by teacher treatment (GLM: χ^2^ = 12.1, *p* = 0.007; Figure 2A). Students exposed to a *M4*-mated teacher selected the *M4* male more frequently than females exposed to no teachers (Model summary: z = 2.3, *p* = 0.019), virgin teachers (Model summary: z = 2.4, *p* = 0.015), and *M7* mated teachers (Model summary: z = 3.3, *p* = 0.001). There was no difference in the preference for *M4* males among the other three treatments (Model summary: z < 1.1, *p* > 0.284). Mate choice of students exposed to *M4*-mated teachers was higher than by random choice (Binomial test: *p* = 0.013), whereas mate choice of students exposed to *M7*-mated teachers was lower than by random choice as less females selected M4 than could be predicted by chance (Binomial test: *p* = 0.048). Mate choice of students exposed to no teacher or a virgin teacher was not different from random choice (Binomial test: *p* > 0.532).

**Figure 2:**
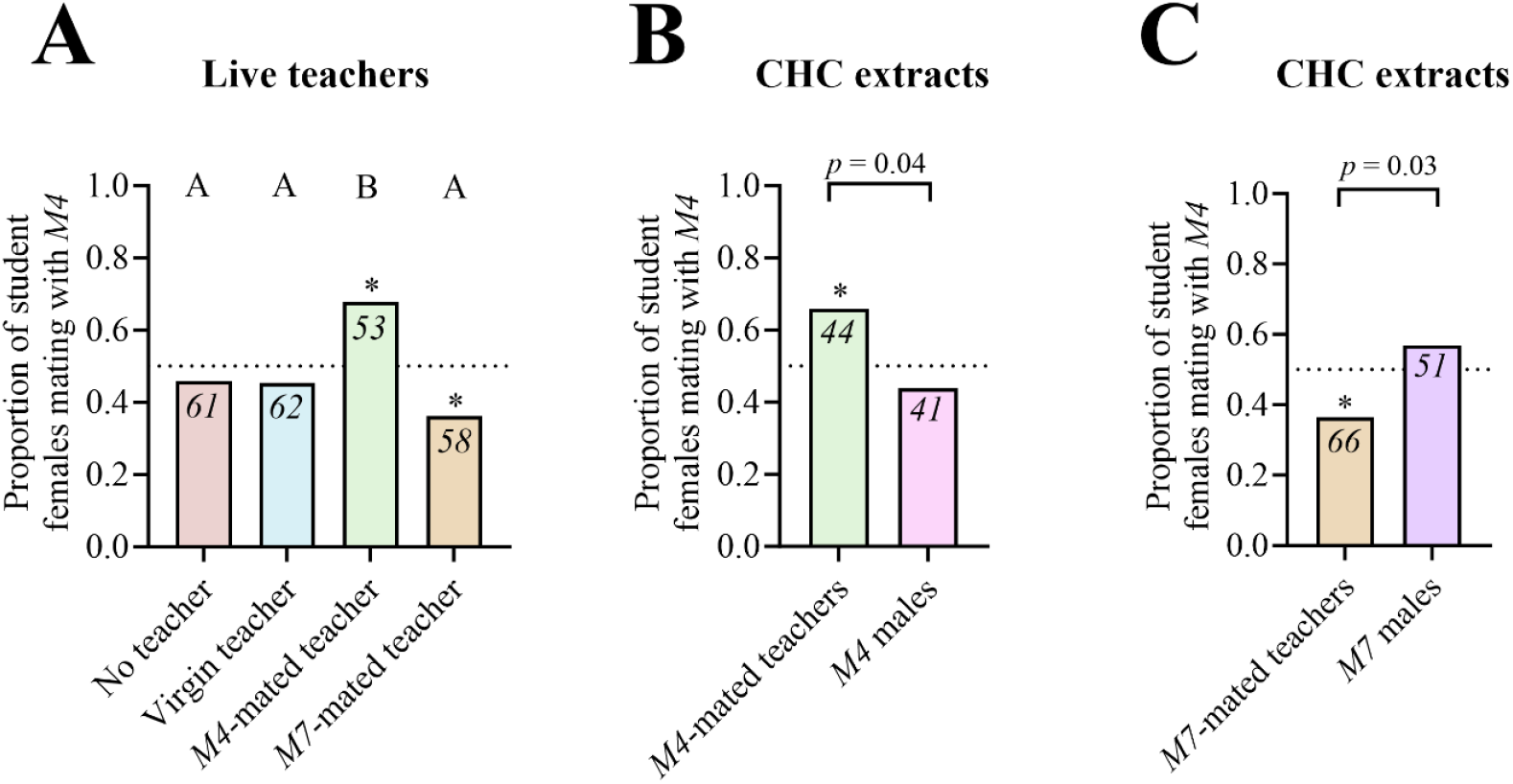
Student preferences for M4 males after teacher and CHC extract exposure. (**A**) Exposure to *M4*-mated teachers increases the preference for *M4* males in student females whereas an exposure to *M7*-mated teachers results in a preference for *M7* males. (**B**) Exposing students to the CHC extracts of *M4*-mated teachers increases their preference for *M4* males, whereas students exposed to the extracts of unmated *M4* males do not exhibit a preference for one of the two males. Bars labelled with the same letters are not statistically different. Numbers in each bar indicate sample size. * above a bar indicates a statistically significant difference from random choice *i*.*e*., 0.5, based on binomial tests. No * indicates no difference from random choice.

Student females exposed to the CHC extracts of *M4*-mated teachers selected the *M4* male more frequently than females exposed to the CHC extracts of *M4* males (GLM: χ^2^ = 4.2, *p* = 0.041; Figure 2B) and selected *M4* males more frequently than predicted by chance (Binomial test: *p* = 0.049) whereas students exposed to extracts of *M4* males displayed no preference for one of the two males (Binomial test: *p* = 0.533). Furthermore, female students exposed to CHC extracts from *M7*-mated teachers chose *M7* males more often than those exposed to CHC extracts from M7 males alone (GLM: χ^2^ = 4.9, *p* = 0.027; Figure 2C). These females also selected *M7* males more frequently than expected by chance (Binomial test: *p* = 0.036), while females exposed only to *M7* male extracts showed no preference between the two males (Binomial test: *p* = 0.401).

The mate choice of students exposed to two teachers was not different from random choice (Binomial test: *p* = 0.542; Figure 3).

**Figure 3:**
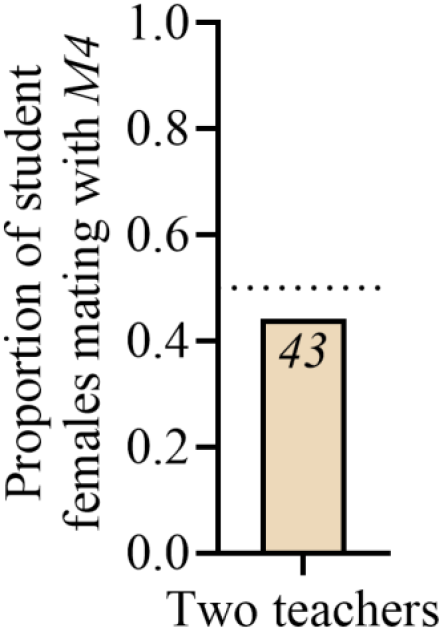
Exposing teachers to two teachers results in lack of preference. Student females were exposed to one M4- and one M7-mated teacher during the teaching phase. No * indicates no difference from random choice *i*.*e*., 0.5, based on binomial tests.

## Discussion

The use of social information during mate choice has been suggested as a fast and low-cost strategy to identify high-quality mating partners (Scauzillo and Ferkin 2019). Most studies on mate choice copying are biased towards vertebrates and visual cues, even if the used species primarily use other sensory modalities, such as olfaction (Scauzillo and Ferkin 2019). In this study, we provide the first evidence of CMCC. Exposure to a recently mated teacher female increases the likelihood that student females will choose a male similar to the one with whom the teacher mated. Student females showed no preference for one of the male genotypes without a teacher, indicating that both male genotypes were largely perceived as comparable in attractiveness, reducing the likelihood that courtship vigor differences alone drove the results. Despite a teacher, some of the student females still chose the other male. Similar observations have been made in studies on visual mate choice copying (e.g., Nöbel et al. 2022*a*), suggesting that mate choice copying may not be absolute. Potential reasons for this incomplete copying might include the social interactions between teacher and student, with students potentially not copying teachers’ decisions when aggressive behaviors were displayed. Copulation duration during teacher mating might further impact the likelihood of CMCC, as a short duration might result in fewer pheromones being transferred to the female. While our findings suggest CMCC as a distinct and novel behavioral phenomenon, we acknowledge that alternative mechanisms, such as associative learning through social experiences with the teacher, could potentially underlie or at least impact the observed preferences. In such a model, females could associate the presented male odor with the positive/negative social experience with the teacher female. Future research investigating these social and behavioral factors will provide valuable insights into how social information spreads through a group and impacts the behavior of other group members.

Mate choice copying via visual cues has previously been reported in *D. melanogaster* (Mery et al. 2009; Dagaeff et al. 2016; Danchin et al. 2018; Nöbel et al. 2018, 2022*b*, 2022*a*, 2023). Our study now provides a novel pathway via which social information can be collected and exploited in the context of mate choice copying. Unlike visual cues, chemical signals enable females to gather social information without directly witnessing mating events, allowing them to assess the mate choices of a broader array of females within their group and to also evaluate mate choice decisions made by other females in the past. The extent to which the neural substrates that integrate visual and chemical cues overlap and how flies respond when these cues provide conflicting information remains to be elucidated. For example, visual mate choice copying requires the expression of the coincidence detector gene *Rutabaga* in the γ-Kanyon cells of the mushroom bodies (Nöbel et al. 2023). These cells receive input not only from visual but also from olfactory sensory neurons and are involved in the regulation of social behavior and responses to social experiences (Sun et al. 2020). It will thus be interesting to study if chemical and visual mate choice copying employs similar neural mechanisms to encode social information and whether γ-Kanyon Cells are involved in this process. Besides mate choice, *D. melanogaster* flies utilize chemical cues provided by the social environment to assess whether another female has chosen strawberries or bananas as an oviposition substrate and to copy this decision (Battesti et al. 2012). Our findings on chemical mate choice copying suggest that the exploitation of social information and copying decisions of other flies via chemical cues may be more prevalent in flies than previously recognized.

The observed behavior in this study also raises broader questions about the mechanisms underpinning social learning and mate choice copying in *D. melanogaster*. One potential mechanism via which mate choice copying can occur is an altered preference for yet-to-be-identified male chemical cues in response to social cues. For example, virgin females are unselective in their initial mating choice due to highly sensitive *OR47b* neurons that sense the male pheromone palmitoleic acid (Kohlmeier et al. 2021). Post-mating desensitization of *OR47b* neurons increases female choosiness as they select only males that produce large amounts of palmitoleic acid for remating. Such studies demonstrate how changes in sensory neuron sensitivity can drive shifts in the preferred male chemotype (Kohlmeier and Billeter, 2023). We now show that exposing virgin females to recently mated females can trigger a preference for specific male pheromone profiles. This social exposure could recalibrate the sensitivity of neurons, either of sensory neurons that sense male pheromones directly or of downstream integrative neurons (Zhang et al. 2021), enhancing responsiveness to particular male pheromone levels. By modifying neural sensitivity to certain male and female pheromones – though the exact compounds remain unidentified – student females may become predisposed to favor males with similar pheromonal profiles to those observed in teacher females.

Most studies investigating visual mate choice copying have employed artificial male phenotypes by marking males with fluorescent powder (Dagaeff et al. 2016; Danchin et al. 2018; Nöbel et al. 2022*a*, 2023). This method complicates the identification of visual cues that females naturally assess, leaving it unclear under which conditions visual mate choice copying occurs (Belkina et al. 2021). On the other hand, many studies investigating male traits assessed by females during mate choice decisions, such as different concentrations of pheromones, often use male genotypes that do not co-occur naturally within the same environment (*e*.*g*., Kohlmeier et al. 2021). This raises the question of whether similar degrees of variation in the investigated male traits occur in nature and, consequently, whether the observed behaviors are ecologically relevant. Our assay involved males from two different isofemale lines, both collected on the same day and in the same environment, ensuring that we examined naturally occurring variation in male traits. By using male genotypes that co-occur and focusing exclusively on their chemical profiles, our approach supports the idea that mate choice copying is an ecologically significant behavior exhibited by females in natural settings.

Our findings reveal that exposure to male extracts alone does not increase student females’ preference for that specific genotype. This indicates that student females must detect cues from both males and mated females simultaneously to enhance their preference for a particular type of male. Functionally, the reliance on cues specific to mated females may increase the likelihood of a female choosing a male already selected by another female, thereby potentially enhancing reproductive success. It is likely that the female cues assessed by the students include compounds only found on females, *i*.*e*., 7,11-Heptacosadiene and 7,11-Nonacosadiene (Billeter and Levine 2013) and/or those compounds characteristic of mated females, such as *cis*-Vaccenyl acetate and 7-Tricosene which are both transferred from the male to the female during copulation (Ejima et al. 2007; Laturney and Billeter 2016). Relying on pheromones characteristic for mated females would additionally protect the female from developing a preference for a specific male in the presence of other virgin females, indicating that this male has not yet been selected by other females. On a mechanistic level, these data suggest that the neuronal circuitry regulating CMCC requires inputs from neurons sensitive to both male and female cues. Which male pheromones are evaluated by the female to identify similar males is unclear. Potential candidates include pheromones previously found to be involved in mate choice, such as Palmitoleic Acid (Lin et al. 2016; Kohlmeier et al. 2021), 7-Tricosene (Grillet et al. 2006), and 7-Pentacosene (Ferveur 2005).

Mate choice copying can increase sexual selection on males, particularly under high-density conditions. Typically, *D. melanogaster* virgin females choose their first mating partner indiscriminately to secure a quick acquisition of sperm. Following the initial mating, they become more selective about male quality to enhance offspring fitness (Kokko and Mappes 2005; Kohlmeier et al. 2021). In scenarios where population density is high and mated females are abundant, mate choice copying can enable virgin females to develop an increased selectiveness for males frequently chosen by others. This allows virgin females to reduce the risk of mating with low-quality males due to the initial lack of choosiness. This mechanism suggests that sexual selection on males is likely to be more intense in populations with high densities. It would be interesting to explore whether the presence of one or a few mated females can initiate a cascade effect among virgin females, leading to a group-level preference for specific male types.

The present data provide initial evidence for CMCC in *D. melanogaster*, yet translating these laboratory findings to the ecology of free-living flies presents intriguing questions. While this study tested student females’ responses to two male genotypes, female flies in the wild may encounter a broader range of males with diverse chemotypes, likely shaped by both genetic and environmental factors. Recent work demonstrated substantial natural variation in CHC profiles across wild populations of *Drosophila*, underscoring the complexity of these profiles in field populations (Ferveur et al. 2024). Currently, however, limited information exists on the population structure of wild *Drosophila*. Seasonal population dynamics, including overwintering females who establish new populations in the summer, add further complexity to our understanding of genetic and chemical diversity in the field. The probability of a student female encountering a male with a chemotype similar to a previously mated “teacher” remains difficult to assess. This is especially challenging as the specific male pheromones driving CMCC have yet to be identified. Though CHC profiles are complex and comprise dozens of compounds (Laturney & Billeter 2016), sexual behaviors and mate choices are driven by few or even single compounds (Billeter and Levine 2013; Laturney and Billeter 2016; Kohlmeier et al. 2021). Therefore, pinpointing the key pheromones involved in CMCC is a crucial next step toward understanding the ecological relevance of this phenomenon in free-living flies.

In summary, we describe chemical mate choice copying as a novel behavior in *D. melanogaster* and our data suggest that female and male cues must be sensed in parallel to induce copying behavior, potentially protecting females from developing a preference for males not previously chosen by other females. This new behavior may impact sexual selection dynamics, particularly when population density is high.

## References

Bateman, A. J. 1948. Intra-sexual selection in Drosophila. Heredity 2:349–368.

Battesti, M., C. Moreno, D. Joly, and F. Mery. 2012. Spread of social information and dynamics of social transmission within Drosophila groups. Current Biology 22:309–313.

Belkina, E. G., A. Shiglik, N. G. Sopilko, S. N. Lysenkov, and A. V. Markov. 2021. Mate choice copying in Drosophila is probably less robust than previously suggested. Animal Behaviour 176:175– 183.

Billeter, J. C., J. Atallah, J. J. Krupp, J. G. Millar, and J. D. Levine. 2009. Specialized cells tag sexual and species identity in Drosophila melanogaster. Nature 461:987–991.

Billeter, J. C., T. P. M. Bailly, and P. Kohlmeier. 2024. The social life of Drosophila melanogaster. Insectes Sociaux, in press.

Billeter, J. C., and J. D. Levine. 2013. Who is he and what is he to you? Recognition in Drosophila melanogaster. Current Opinion in Neurobiology 23:17–23.

Champagnon, J., and R. Cueva del Castillo. 2008. Female mate choice, calling song and genetic variance in the cricket, Gryllodes sigillatus. Ethology 114:223–230.

Claudio-Piedras, F., B. Recio-Tótoro, J. Cime-Castillo, R. Condé, M. Maffei, and H. Lanz-Mendoza. 2021. Dietary and Plasmodium challenge effects on the cuticular hydrocarbon profile of Anopheles albimanus. Scientific Reports 11:11258.

Dagaeff, A. C., A. Pocheville, S. Nöbel, A. Loyau, G. Isabel, and E. Danchin. 2016. Drosophila mate copying correlates with atmospheric pressure in a speed learning situation. Animal Behaviour 121:163–174.

Danchin, E., S. Nöbel, A. Pocheville, A.-C. Dagaeff, L. Demay, M. Alphand, S. Ranty-Roby, et al. 2018. Cultural flies: Conformist social learning in fruitflies predicts long-lasting mate-choice traditions. Science 362:1025–1030.

Doubovetzky, N., P. Kohlmeier, S. Bal, and J.-C. Billeter. 2024. Cryptic female choice in response to male pheromones in Drosophila melanogaster. Current Biology 34:4539–4546.

Drullion, D., and F. Dubois. 2008. Mate-choice copying by female zebra finches, Taeniopygia guttata: What happens when model females provide inconsistent information? Behavioral Ecology and Sociobiology 63:269–276.

Dugatkin, L. A., and J.-G. J. Godin. 1993. Female mate copying in the guppy (Poecilia reticulata): age-dependent effects. Behavioral Ecology 4:289–292.

Ejima, A., B. P. C. Smith, C. Lucas, W. Van Der Goes Van Naters, C. J. Miller, J. R. Carlson, J. D. Levine, et al. 2007. Generalization of courtship learning in Drosophila is mediated by cis-vaccenyl acetate. Current Biology 17:599–605.

Everaerts, C., J. P. Farine, M. Cobb, and J. F. Ferveur. 2010. Drosophila cuticular hydrocarbons revisited: Mating status alters cuticular profiles. PLoS ONE 5:e9607.

Ferveur, J. F. 2005. Cuticular hydrocarbons: Their evolution and roles in Drosophila pheromonal communication. Behavior Genetics 35:279–295.

Ferveur, J. F., J. Cortot, M. Cobb, and C. Everaerts. 2024. Natural diversity of cuticular pheromones in a local population of Drosophila after laboratory acclimation. Insects 15:273.

Fricke, C., A. Bretman, and T. Chapman. 2008. Adult male nutrition and reproductive success in Drosophila melanogaster. Evolution 62:3170–3177.

Grillet, M., L. Dartevelle, and J. F. Ferveur. 2006. A Drosophila male pheromone affects female sexual receptivity. Proceedings of the Royal Society B: Biological Sciences 273:315–323.

Hill, G. E. 2002. A red bird in a brown bag. Oxford University Press, New York.

Jallon, J. M. 1984. A few chemical words exchanged by Drosophila during courtship and mating. Behavior Genetics 14:441–478.

Kavaliers, M., R. Matta, and E. Choleris. 2017. Mate-choice copying, social information processing, and the roles of oxytocin. Neuroscience and Biobehavioral Reviews 72:232–242.

Kohlmeier, P., and J.-C. Billeter. 2023. Genetic mechanisms modulating behaviour through plastic chemosensory responses in insects. Molecular Ecology 32:45–60.

Kohlmeier, P., Y. Zhang, J. Gortner, C.-Y. Su, and J.-C. Billeter. 2021. Mating increases Drosophila melanogaster females’ choosiness by reducing olfactory sensitivity to a male pheromone. Nature Ecology & Evolution 5:1165–1173.

Kokko, H., and J. Mappes. 2005. Sexual selection when fertilization is not guaranteed. Evolution 59:1876–1885.

Kurtovic, A., A. Widmer, and B. J. Dickson. 2007. A single class of olfactory neurons mediates behavioural responses to a Drosophila sex pheromone. Nature 446:542–546.

Laturney, M., and J.-C. Billeter. 2016. Drosophila melanogaster females restore their attractiveness after mating by removing male anti-aphrodisiac pheromones. Nature Communications 7:12322.

Lin, H. H., D. S. Cao, S. Sethi, Z. Zeng, J. S. R. Chin, T. S. Chakraborty, A. K. Shepherd, et al. 2016. Hormonal modulation of pheromone detection enhances male courtship success. Neuron 90:1272– 1285.

Lung, O., and M. F. Wolfner. 2001. Identification and characterization of the major Drosophila melanogaster mating plug protein. Insect Biochemistry and Molecular Biology 31:543–551.

Mane, S. D., L. Tompkins, and R. C. Richmond. 1983. Male Esterase 6 catalyzes the synthesis of a sex pheromone in Drosophila melanogaster Females. Science 222:419–421.

Markow, T. A. 1988. Reproductive behavior of Drosophila melanogaster and D. nigrospiracula in the field and in the laboratory. Journal of Comparative Psychology 102:169–173.

Mery, F., S. A. M. Varela, É. Danchin, S. Blanchet, D. Parejo, I. Coolen, and R. H. Wagner. 2009. Public versus personal information for mate copying in an invertebrate. Current Biology 19:730–734.

Milinski, M., and T. C. M. Bakker. 1992. Costs influences sequential mate choice in sticklebacks, Gasterosteus aculeatus. Proceedings of the Royal Society of London. Series B: Biological Sciences 250:229–233.

Møller, A. P., and A. Pomiankowski. 1993. Fluctuating asymmetry and sexual selection. Genetica 89:267–279.

Muria, A., P. Y. Musso, M. Durrieu, F. R. Portugal, B. Ronsin, M. D. Gordon, R. Jeanson, et al. 2021. Social facilitation of long-lasting memory is mediated by CO2 in Drosophila. Current Biology 31:2065–2074.

Nöbel, S., E. Danchin, and G. Isabel. 2018. Mate-copying for a costly variant in Drosophila melanogaster females. Behavioral Ecology 29:1150–1156.

Nöbel, S., E. Danchin., and G. Isabel. 2023. Mate copying requires the coincidence detector Rutabaga in the mushroom bodies of Drosophila melanogaster. iScience 26:107682.

Nöbel, S., M. Monier, L. Fargeot, G. Lespagnol, E. Danchin, and G. Isabel. 2022a. Female fruit flies copy the acceptance, but not the rejection, of a mate. Behavioral Ecology 33:1018–1024.

Nöbel, S., M. Monier, D. Villa, É. Danchin, and G. Isabel. 2022b. 2-D sex images elicit mate copying in fruit flies. Scientific Reports 12:22127.

Scauzillo, R. C., and M. H. Ferkin. 2019. Factors that affect non-independent mate choice. Biological Journal of the Linnean Society 128:499–514.

Sun, Y., R. Qiu, X. Li, Y. Cheng, S. Gao, F. Kong, L. Liu, et al. 2020. Social attraction in Drosophila is regulated by the mushroom body and serotonergic system. Nature Communications 11:5350.

Sureau, G., and J. F. Ferveur. 1999. Co-adaptation of pheromone production and behavioural responses in Drosophila melanogaster males. Genetical Research 74:129–137.

Trivers, R. L. 1972. Parental investment and sexual selection. Pages 52–95 in Sexual Selection and the Descent of Man: The Darwinian Pivot. Aldine, Chicago.

Verschut, T. A., P. Kohlmeier, and J.-C. Billeter. 2022. Bioassaying the function of pheromones in Drosophila melanogaster’s social behavior. Pages 123–157 in Behavioral Neurogenetics.

Verschut, T. A., R. Ng, N. P. Doubovetzky, G. Le Calvez, J. L. Sneep, A. J. Minnaard, C.-Y. Su, et al. 2023. Aggregation pheromones have a non-linear effect on oviposition behavior in Drosophila melanogaster. Nature Communications 14:1544.

Villella, A., and J. C. Hall. 2008. Chapter 3 Neurogenetics of courtship and mating in Drosophila. Pages 67–184 in Advances in Genetics.

Witte, K. 2007. Learning and Mate Choice. Pages 70–95 in Fish Cognition and Behavior. Wiley Blackwell.

Wyatt, T. D. 2003. Pheromones and animal behaviour (1st ed.). Cambridge University Press, Cambridge.

Zawistowski, S., and R. C. Richmond. 1986. Inhibition of courtship and mating of Drosophila melanogaster by the male-produced lipid, cis-Vaccenyl Acetate. Journal of Insect Physiology 32:189– 192.

Zhang, S., E. Glantz, D. Rogulja, and M. Crickmore. 2021. Hormonal control of motivational circuitry orchestrates the transition to sexuality in Drosophila. Science Advances 7:eabg6926.

